# High-resolution genetic map and QTL analysis of growth-related traits of *Hevea brasiliensis* cultivated under suboptimal temperature and humidity conditions

**DOI:** 10.1101/355008

**Authors:** André Ricardo Oliveira Conson, Cristiane Hayumi Taniguti, Rodrigo Rampazo Amadeu, Isabela Aparecida Araújo Andreotti, Livia Moura de Souza, Luciano Henrique Braz dos Santos, João Ricardo Bachega Feijó Rosa, Camila Campos Mantello, Carla Cristina da Silva, Erivaldo José Scaloppi Junior, Rafael Vasconcelos Ribeiro, Vincent Le Guen, Antonio Augusto Franco Garcia, Paulo de Souza Gonçalves, Anete Pereira de Souza

## Abstract

Rubber tree (*Hevea brasiliensis*) cultivation is the main source of natural rubber worldwide and has been extended to areas with suboptimal climates and lengthy drought periods; this transition affects growth and latex production. High-density genetic maps with reliable markers support precise mapping of quantitative trait loci (QTL), which can help reveal the complex genome of the species, provide tools to enhance molecular breeding, and shorten the breeding cycle. In this study, QTL mapping of the stem diameter, tree height, and number of whorls was performed for a full-sibling population derived from a GT1 and RRIM701 cross. A total of 225 simple sequence repeats (SSRs) and 186 single-nucleotide polymorphism (SNP) markers were used to construct a base map with 18 linkage groups and to anchor 671 SNPs from genotyping by sequencing (GBS) to produce a very dense linkage map with small intervals between loci. The final map was composed of 1,079 markers, spanned 3,779.7 cM with an average marker density of 3.5 cM, and showed collinearity between markers from previous studies. Significant variation in phenotypic characteristics was found over a 59-month evaluation period with a total of 38 QTLs being identified through a composite interval mapping method. Linkage group 4 showed the greatest number of QTLs (7), with phenotypic explained values varying from 7.67% to 14.07%. Additionally, we estimated segregation patterns, dominance, and additive effects for each QTL. A total of 53 significant effects for stem diameter were observed, and these effects were mostly related to additivity in the GT1 clone. Associating accurate genome assemblies and genetic maps represents a promising strategy for identifying the genetic basis of phenotypic traits in rubber trees. Then, further research can benefit from the QTLs identified herein, providing a better understanding of the key determinant genes associated with growth of *Hevea brasiliensis* under limiting water conditions.

## Introduction

*Hevea brasiliensis* (Willd. ex. Adr. de Juss.) Muell-Arg., also known as the rubber tree, is one of the most symbolic species from the Amazon basin owing to its worldwide use in natural latex production, the global demand for which suggests expansion of the industry (Warren-Thomas *et al*., 2015). It is a perennial and allogamous tree from the Euphorbiaceae family. Its breeding has started in 1876 (Priyadarshan and Clément-Demange, 2004). The main trait in domestication of this tree is yield and its improvement along the twentieth century is related with factors such as growth of the trunk during immature phase (Priyadarshan, 2017). Girth measurements has been the best and most practicable measure to determine the criterion of tappability along the tree cycle (Dijkman, 1951).

The species contains 36 chromosomes (2n = 36), and studies indicate that *H. brasiliensis* is an amphidiploid that has been stabilized its genome during the course of evolution (Saha and Priyadarshan, 2012). Because of its genome size and high repeat content (Lau *et al*., 2016; Tang *et al*., 2016), genetic analysis of the species represents considerable obstacles to breeding, and several methodologies must be used to better understand its organization (Fierst, 2015).

Rubber tree plantations are mainly located in Southeast Asia under optimal edaphoclimatic conditions due to the occurrence of the South America Leaf Blight (SALB) disease in Latin America. Because the species can be cultivated under suboptimal conditions, plantations in “escape areas”, such as the southeastern region of Brazil, where epidemic SALB has not been found, might be an alternative for natural rubber production (Gonçalves *et al*., 2011; Souza *et al*., 2013; Jaimes *et al*., 2016). However, suboptimal conditions (temperature and humidity) affect plant development, consequently affecting rubber production (Silpi *et al*., 2006; Rao and Kole, 2016).

Developing new rubber tree clones may extend up to 20–30 years and becomes expensive. Molecular markers can help identify allelic variants related to phenotypes adapted to these regions and can shorten the breeding cycle (Priyadarshan and Clément-Demange, 2004). Therefore, significant results obtained for other crops using effective and cost-efficient microsatellite markers associated with next-generation sequencing (NGS) technologies, such as the simultaneous discovery and genotyping of thousands of single-nucleotide polymorphisms (SNPs) through genotyping by sequencing (GBS) (Elshire *et al*., 2011), can be very useful for *H. brasiliensis*.

Because no inbred lines are available for the rubber tree species, the mapping methods for these species must be adapted to full-sibling populations, once there are more alleles involved and linkage phases to be estimated compared with the population from inbred lines. The first and most widespread method for building genetic maps for outcrossing species is the pseudo-testcross, which allowed the first genetic mapping studies in such species, but it considers only markers segregating 1:1 and results in separated maps for each parent (Grattapaglia and Sederoff, 1994). Later, several methods were proposed, including maximum likelihood estimators for all types of markers segregation (Maliepaard et al., 1997) and the possibility to build integrated genetic maps using information from all available markers with multipoint estimation (Wu *et al*., 2002b, 2002a). In the rubber tree, different methods have been used to obtain higher-resolution genetic maps (Pootakham *et al*., 2015; Shearman *et al*., 2015) than those obtained in previous studies (Lespinasse *et al*., 2000; Souza *et al*., 2013). Methods with higher-coverage genome reference sequences of the species are also important to detect markers associated with quantitative trait loci (QTL).

In addition to a reliable map, in QTL studies, it is also important to use a robust experimental design. Forestry experiments require large field areas due to the needed spacing for each tree. Therefore, the field area size limits the number of possible genotype replications in the design. Augmented blocks are a very efficient and widely used design proposed to address reduced numbers of replicates (Federer and Raghavarao, 1975). Joint analysis is possible using information from checks included in all incomplete blocks. To improve the statistical power in an augmented blocks design, a possible approach is to use replicated augmented blocks. After modeling and evaluating the experiment, it is possible to gather the genetic values for each genotype and to use them alongside the map in QTL mapping methods. The QTL method must also be adapted to outcrossing species. Gazaffi et al., (2014) developed a composite interval mapping (CIM) model for outcrossing species considering integrated maps. The model uses maximum likelihood and mixture models to estimate three orthogonal contrasts, which are used to estimate additive and dominance effects and QTL segregation patterns and linkage phases within the molecular markers. The QTL screening includes cofactors to avoid influences from QTLs outside the mapping interval.

Here, we report an integrated high-density genetic map for *Hevea brasiliensis* obtained from a set of high-quality SNP and simple sequence repeat (SSR) markers. Moreover, the experiment was conducted in suboptimal environment, in order to explore QTLs under this condition. QTL mapping for complex traits related to plant growth (stem diameter, tree height, and the number of whorls) was performed over a 59-month evaluation period, allowing for more accurate QTL localization.

## Materials and methods

### Mapping population and DNA extraction

An F1 population containing 146 individuals was derived from open pollination between the genotypes GT1 and RRIM701. The GT1 clone, which has been widely used in *Hevea* breeding since the 1920s, is classified as a primary clone that is male sterile (Shearman *et al*., 2015) and tolerant to wind and cold. From the Asian breeding program, RRIM701 exhibits vigorous growth and an increased circumference after initial tapping (Romain and Thierry, 2011).

Effective pollination was confirmed using 10 microsatellite markers that were selected based on the polymorphism information content (PIC) and the polymorphisms between the parents. After removing two contaminants, the mapping population was planted at the Center of Rubber Tree and Agroforestry Systems/Agronomic Institute (IAC) in the northwest region of São Paulo state (20º 25’ 00” S and 49º 59’ 00” W at a 450-meter altitude), Brazil, in 2012. The climate is the Köppen Aw type (Alvares *et al*., 2013). Based on air temperature and rainfall, the water balance was calculated using the method developed by Thornthwaite and Mather (1955), adopting 100 mm as the maximum soil water holding capacity. An augmented block design was used (Federer and Raghavarao, 1975). To improve accuracy, the trial (one augmented block design) was repeated side-by-side four times with different randomizations. Each trial has four blocks and two plants (clones) per plot spaced at 4 m by 4 m. Hereafter, trial is considered a repetition. Therefore, each block consisted of 37 genotypes of GT1 x RRIM701 crosses and four checks (GT1, PB235, RRIM701 and RRIM600). The experiment has a total of 656 (41 plots x 4 blocks x 4 repetition) plots and 1,312 trees.

Genomic DNA was extracted from lyophilized leaves using 2% cetyltrimethylammonium bromide (CTAB) as described by Doyle and Doyle (1987). DNA integrity was assessed by electrophoresis on an ethidium bromide-stained 1% agarose gel, and its concentration was estimated with a QuantiFluor-ST fluorometer (Promega, Madison, WI, USA).

### Microsatellite analysis

A total of 1,190 SSR markers, comprising 476 genomic markers (Souza *et al*., 2009; Le Guen *et al*., 2011; Mantello *et al*., 2012), 353 expressed sequence tag (EST)-SSR markers (Feng *et al*., 2009; Silva *et al*., 2014), and 361 additional unpublished EST-SSRs identified in cDNA libraries constructed from subtractive suppression hybridization libraries (Garcia *et al*., 2011), were tested to verify polymorphisms between the parents. Subsequently, from 569 (48%) polymorphic markers, those with a more informative segregation pattern (Wu et al., 2002b) and with a clear profile were selected, totaling 287 SSRs for the population genotyping.

The amplification of SSRs was performed as previously described by the authors. PCR products were visualized using three different techniques: (1) silver nitrate staining (Creste *et al*., 2001); (2) LI-COR 4300 DNA analyzer (LI-COR Inc., Lincoln, NE, USA), and (3) ABI 3500XL DNA sequencer (Life Technologies, Carlsbad, CA, USA). Markers that were considered monomorphic or homozygous for both parents and markers without clear bands were excluded.

### SNP analysis

SNP markers were developed from full-length EST libraries (Silva *et al*., 2014) and a bark transcriptome (Mantello *et al*., 2014). A total of 391 SNPs was selected for genotyping using the Sequenom MassARRAY Assay Design program (Agena Bioscience^®^, San Diego, CA, USA) with the following parameters: (1) a high plex preset with a multiplex level = 24, (2) an amplicon length varying from 80 to 200 bp, and (3) number of iterations = 10. SNPs were selected for population genotyping using the Sequenom iPLEX MassARRAY^®^ (Sequenom Inc., San Diego, California, USA) and were annotated within the mevalonate and 2-C-methyl-D-erythritol 4-phosphate (MEP) pathways, which are involved in latex biosynthesis and wood synthesis (Table S1). The first amplification reaction was performed using 1–3 ng of genomic DNA, and the subsequent steps were performed following the manufacturer’s instructions. MassArray Typer v4.0 software (Sequenom, Inc.) was used for allele visualization.

Ninety-six other polymorphic SNPs obtained from genomic (Souza *et al*., 2016) and transcriptomic (Salgado *et al*., 2014) data were also selected for genotyping using the BioMark™ Real-Time PCR system with the 96.96 Dynamic Array™ IFC (Fluidigm Corporation, San Francisco, CA, USA). For kompetitive allele-specific PCR (KASP) assays, 1.4 μL of KASP assay reagent, 5 μL of 2X assay loading reagent, 3.6 μL of ultrapure water, and 2.8 μL of KASP assay mix (12 μM allele-specific forward primer and 30 μM reverse primer) were used. Furthermore, 4.73 μL of KASP 2X reagent mix, 0.47 μL of 20X GT sample loading reagent, 0.3 μL of ultrapure water, and 240 ng of DNA were used for PCR. The thermal conditions for fragment amplification were 94°C for 15 minutes; 94°C for 10 seconds; 57°C for 10 seconds; 72°C for 10 seconds; 10 cycles of 94°C for 20 seconds and 65°C to 57°C for 1 minute (−0.8°C per cycle); and 26 cycles of 94°C for 20 seconds and 57°C for 1 minute. The genotyping data were analyzed using the BioMark™ SNP Genotyping Analysis v2.1.1 program.

To saturate the linkage map with more SNPs, we performed the GBS protocol as described by Elshire *et al*. (2011). GBS libraries were generated from 100 ng of genomic DNA, and the *Eco*T22I enzyme was used to reduce the genomic complexity. Library quality checks were performed using the Agilent DNA 1000 Kit with the Agilent 2100 Bioanalyzer (Agilent Technologies, Santa Clara, USA). Library sequencing was performed on an Illumina GAIIx platform (Illumina Inc., San Diego, CA, USA). Quality assessment of the generated sequence data was performed with the NGS QC Toolkit v2.3.3 (Patel and Jain, 2012).

Sequencing data analysis was performed using TASSEL-GBS v3.0 (Bradbury *et al*., 2007). Retained tags with a minimum count of six reads were aligned to the *H. brasiliensis* reference genome sequence, GenBank: LVXX00000000.1 (Tang et al., 2016), using Bowtie2 version 2.1 (Langmead and Salzberg, 2012). The obtained variant call format (VCF) files were filtered using VCFtools (Danecek *et al*., 2011), and only biallelic SNPs, with a maximum of 25% missing data, were retained. Additionally, SNPs were filtered by removing redundant markers, segregation-distorted markers, and markers homozygous for both parents using the R package OneMap v2.1.1 (Margarido *et al*., 2007).

### Genetic mapping

An integrated linkage map was developed in two stages in which all markers were tested for segregation distortion (p-value < 0.05) with Bonferroni’s multiple test correction. All of the following logarithm of the odds (LOD) thresholds were also obtained based on Bonferroni’s multiple test correction according to the number of two point tests. First, a base map with SSR and SNP markers (except GBS-based SNPs) was built, and the markers were grouped using an LOD value of 5.37 and a maximum recombination frequency of 0.4. The markers were ordered through a hidden Markov chain multipoint approach (Lander and Green, 1987) considering outcrossing species (Wu *et al*., 2002a). The most informative markers (1:1:1:1) were ordered using exhaustive search, and the remaining markers were positioned using the TRY algorithm (Lander and Green, 1987). The RIPPLE algorithm (Lander and Green, 1987) was used to improve final orders. The marker order of this first map was fixed before insertion of the GBS-based markers. Later, the GBS markers were grouped into the base map considering an LOD of 7.22 and a maximum recombination fraction of 0.4. The best position for each marker was identified using the TRY and RIPPLE algorithms. All of the analyses were performed using OneMap software v2.1.1 (Margarido *et al*., 2007; Mollinari *et al*., 2009). The Kosambi mapping function was used (Kosambi, 1943) to estimate map distances.

### Phenotypic analysis

Tree height, number of whorls, and stem diameter at 50 cm above ground level were measured to evaluate the growth of individual trees, with mean values calculated based on two plants per plot. Because the early attainment of tappable girth is one of the main selection traits for *H. brasiliensis* breeding (Rao and Kole, 2016), stem diameters were measured at seven different ages between July 2013 and June 2017 (12, 17, 22, 28, 35, 47, and 59 months). The tree heights and numbers of whorls were measured at 17 and 22 months.

The statistical linear mixed model used for the analysis was as follows:

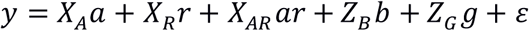
where *y* is the vector with the mean phenotypic trait per plot; and *X_A_*, *X_R_*, and *X_AR_* are the incidence matrices for the fixed effects for age (*a*), replicate (*r*), and interaction age *x* replicate (*ar*). *Z_B_* and *Z_G_* are the incidence matrices for the random effects for block (*b*) and genotype (*g*), with *b* ~ *MVN*(0,*I_B_* ⊗ *I_R_* ⊗ *G_AB_*) and *g* ~ *MVN*(0,*I_G_* ⊗ *G_AG_*), where *I_n_* is an identity matrix with order *n*; ⊗ is the Kronecker product; *MVN* stands for Multivariate Normal Distribution; and *G_AB_* and *G_AG_* are the variance-covariance matrices for the effects of replicates within blocks within age and within genotype within age, respectively. *ε* is the residual, where *ε* ~ *MVN*(0,*R_c_* ⊗ *R_R_* ⊗ *R_A_*), where *R_c_*, *R_R_*, and *R_A_* are the variance-covariance matrices for the column, row, and age for the residual effects, respectively. Different variance-covariance structures for *V_AB_*, *V_AG_*, *R_c_*, *R_R_*, and *R_A_* were adjusted and compared based on the Akaike information criterion (AIC) (Akaike, 1974) and the Bayesian information criterion (BIC) (Schwarz, 1978) values (Table S2). The variance components were estimated by maximizing the maximum likelihood distribution (REML). Analyses were performed using the packages ASReml-R v3 (Gilmour *et al*., 2009) and ASRemlPlus (Brien, 2016). The heritability (*H*^2^) for each age was estimated as presented by Cullis *et al*. (2006) and Piepho and Möhring (2007), where 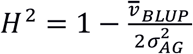 and 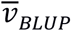 indicates the mean variance between the two best linear unbiased predictions (BLUPs). After the phenotypic analysis, we used the differences between estimated BLUPs of subsequent ages for QTL mapping. As an example, for a genotype with a BLUP of 10 cm of diameter at 17 months and with a BLUP of 13 cm of diameter at 22 months, the difference of BLUPs (i.e., 3 cm) was used in the QTL mapping.

### QTL mapping

QTL mapping for stem diameter, tree height, and number of whorls traits was performed by applying a CIM model to full-sib families (Gazaffi *et al*., 2014). This method uses a multipoint approach that considers outcrossing segregation patterns and, through contrasts, estimates three genetic effects, one for each parent (hereafter called the additive effect) and one for their interaction (hereafter called the dominance effect). For the CIM model, QTL genotype probabilities were obtained for each 1-cM interval considering a window size of 15 cM. Cofactors were identified for each linkage group using stepwise selection based on the AIC (Akaike, 1974) to select the models. Furthermore, a maximum number of parameters, 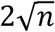, where *n* is the number of individuals, was used to avoid super-parametrization (Gazaffi *et al*., 2014). Subsequently, only effects with a 5% significance level were retained in the cofactor model. QTLs were defined according to a threshold based on 5% significance across distributed LOD scores obtained by selecting the second LOD profile peak from 1,000 permutations (Chen and Storey, 2006). Additionally, for each CIM model, the additive genetic effects for each parent, the dominance effects, and the linkage phases, segregation patterns, and proportions of phenotypic variations explained by the QTLs (R2) were estimated. Analyses were performed using the R package fullsibQTL (Gazaffi *et al*., 2014).

## Results

### Filtering and polymorphism analysis

For the linkages analysis 239 SNP markers from genomic, transcriptomic, and cDNA datasets were retained. The marker segregation for the SSRs and SNPs are shown in Table 1. All genotyped markers, including 33 markers (6.3%) identified with distortion from the expected Mendelian segregation ratio, were retained in the linkage analysis.

**Table 1.**
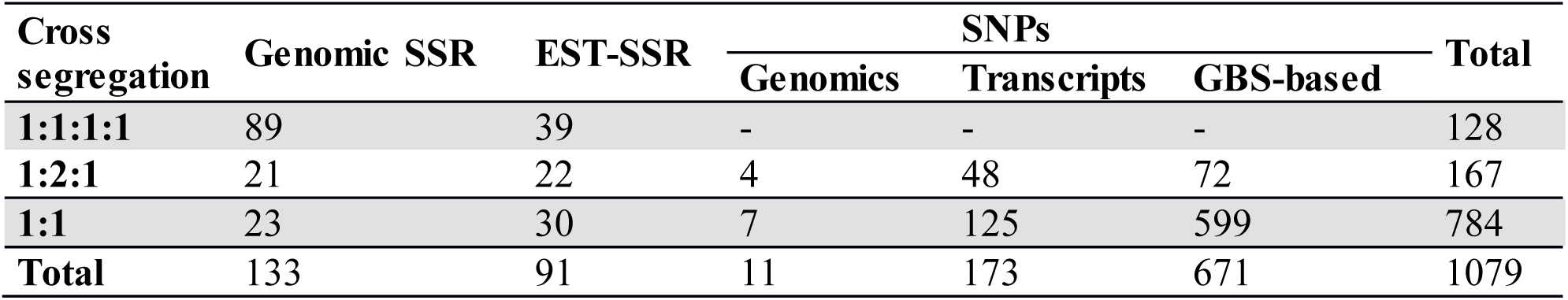
Overall segregation of SSR and SNP markers mapped in the final map for progenies of the GT1 and RRIM701 cross.

The GBS library generated a total of 68,777,856 raw reads (90.60% Q20 bases), with an average of 474,330 reads per individual. After alignment to the *H. brasiliensis* reference genome, 81,832 SNPs were identified. In total, 53,836 markers with more than 25% missing data and/or a non-biallelic status, 1,587 redundant markers, 17,865 markers with two homozygous parents, and 5,546 markers with segregation distortion were eliminated. Thus, a total of 2,998 (3.66% of the initial set of markers) high-quality GBS-based SNPs were retained, most of which were classified in a 1:1 fashion (2,371 SNPs).

### Genetic mapping

A total of 411 markers were mapped in the base map constructed from a set of 526 markers (287 SSRs and 239 SNPs). The markers were distributed over 18 linkage groups (LG) in 2,482.3 cM (Figure 1), and the length of each group ranged from 78.3 cM (LG4) to 191.9 cM (LG14). Linkage group 10 showed the highest average density and the highest number of markers mapped (53), whereas linkage group 15 showed the lowest density (Table 2).

**Table 2.** Linkage group information regarding the base and final maps from the cross between GT1 and RRIM701. LG – linkage group; length - mean length in cM; density - mean density in cM; gap - length of larger gap in cM.

| LG | Base Map |  |  |  |  | Final Map |  |  |  |  |
| --- | --- | --- | --- | --- | --- | --- | --- | --- | --- | --- |
|  | SSRs | SNPs | Length (cM) | Density (cM) | Gap (cM) | SSRs | SNPs | Length (cM) | Density (cM) | Gap (cM) |
| **1** | 14 | 17 | 153.1 | 4.9 | 18.4 | 13 | 53 | 232.4 | 3.5 | 13.0 |
| **2** | 19 | 8 | 140.2 | 5.2 | 14.4 | 19 | 33 | 199.3 | 3.9 | 9.1 |
| **3** | 14 | 6 | 115.0 | 5.8 | 20.7 | 14 | 43 | 184.9 | 3.2 | 14.0 |
| **4** | 6 | 5 | 78.3 | 7.1 | 17.9 | 6 | 39 | 163.8 | 3.6 | 12.9 |
| **5** | 9 | 13 | 176.9 | 8.0 | 24.9 | 9 | 65 | 272.0 | 3.7 | 16.6 |
| **6** | 17 | 14 | 162.9 | 5.3 | 20.1 | 17 | 63 | 228.1 | 2.9 | 11.6 |
| **7** | 12 | 5 | 80.3 | 4.7 | 19.4 | 12 | 32 | 109.4 | 2.5 | 14.1 |
| **8** | 14 | 8 | 141.3 | 6.4 | 18.4 | 14 | 45 | 229.7 | 3.9 | 18.5 |
| **9** | 13 | 6 | 124.6 | 6.6 | 20.8 | 13 | 33 | 196.3 | 4.3 | 14.3 |
| **10** | 26 | 27 | 185.4 | 3.5 | 16.2 | 26 | 52 | 228.1 | 2.9 | 11.5 |
| **11** | 6 | 15 | 164.8 | 7.8 | 27.5 | 6 | 58 | 235.4 | 3.7 | 20.6 |
| **12** | 7 | 10 | 104.1 | 6.1 | 23.3 | 7 | 32 | 167.5 | 4.3 | 12.8 |
| **13** | 6 | 10 | 136.7 | 8.5 | 24.2 | 6 | 58 | 218.7 | 3.4 | 15.0 |
| **14** | 22 | 13 | 191.9 | 5.5 | 15.6 | 22 | 51 | 258.8 | 3.5 | 17.0 |
| **15** | 10 | 6 | 160.0 | 10.0 | 32.5 | 10 | 39 | 225.3 | 4.6 | 16.2 |
| **16** | 10 | 6 | 100.8 | 6.3 | 17.3 | 10 | 53 | 166.0 | 2.6 | 12.9 |
| **17** | 6 | 7 | 107.5 | 8.3 | 19.6 | 6 | 54 | 206.7 | 3.6 | 9.8 |
| **18** | 14 | 10 | 158.5 | 6.6 | 26.0 | 14 | 52 | 257.3 | 3.9 | 17.1 |
| **Total** | 225 | 186 | 2482.3 | 6.0 | 32.5 | 224 | 855 | 3779.7 | 3.5 | 20.6 |

**Figure 1.**
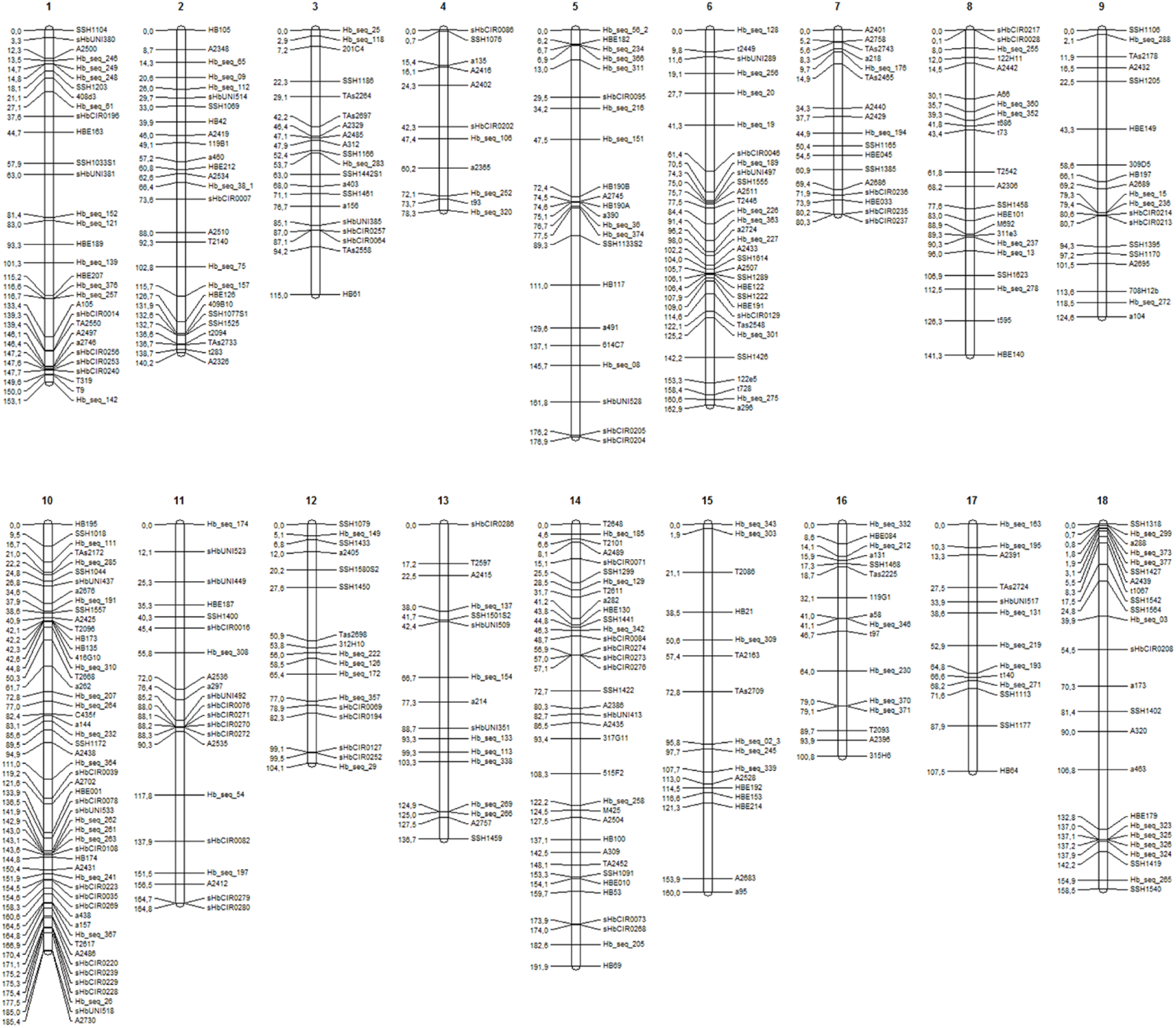
LGs (1 to 18) for the base genetic map, including 411 markers (225 SSRs and 186 SNPs).

The final genetic map, constructed using advances from the previous base map and the numerous high-quality SNPs produced with the GBS technique, was based on 408 previously mapped markers and 671 SNPs identified from GBS (Bioproject PRJEB25899). One SSR and two SNP markers were removed from the base map to build the final map because there were inconsistencies in their primers. Therefore, 1,079 mapped markers were distributed over 18 LGs and covered a total of 3,779.7 cM (Figure 2). The length of each group ranged from 109.4 cM and 44 markers (LG7) to 272 cM and 74 markers (LG5), with average marker densities of 2.5 and 3.7 cM, respectively.

**Figure 2.**
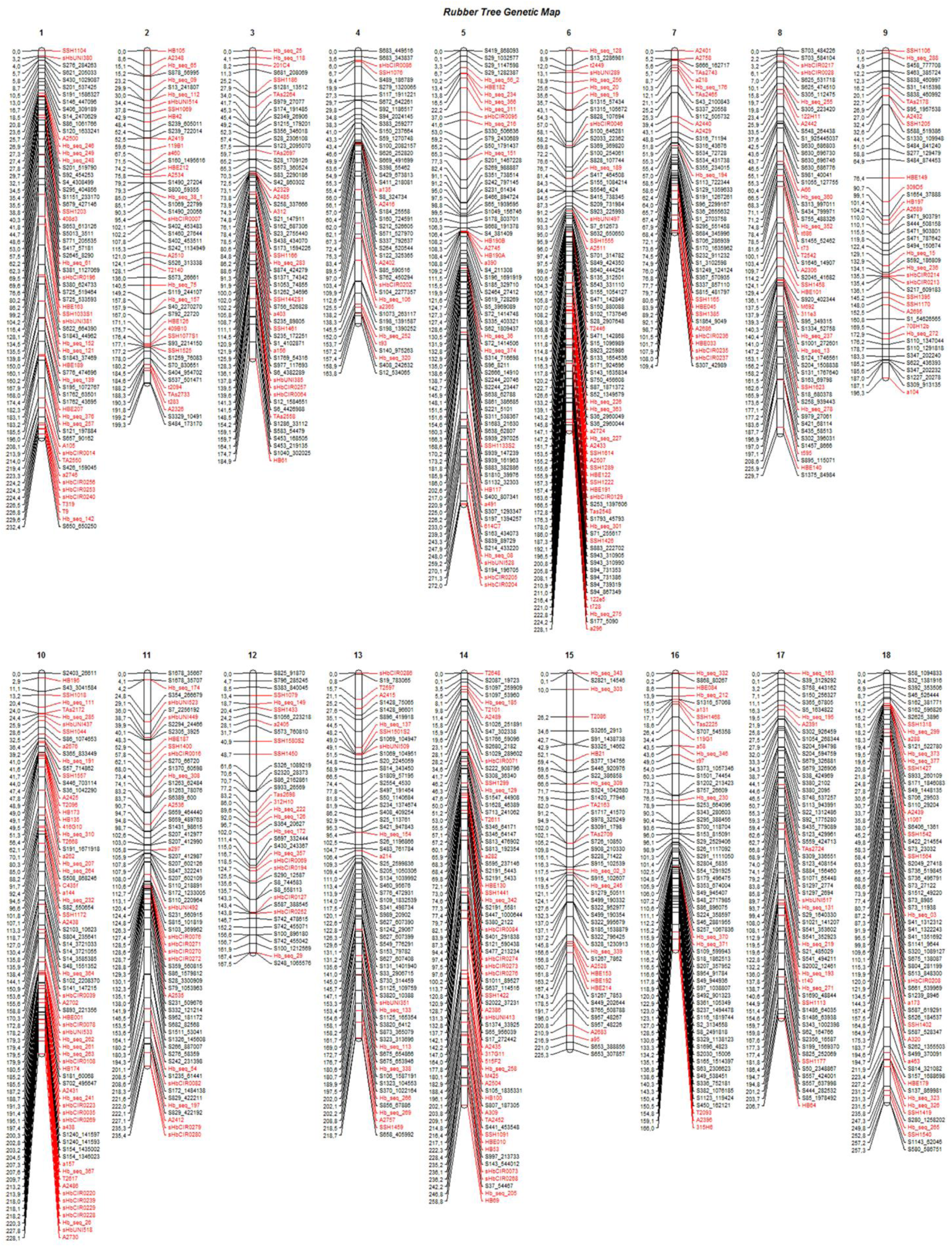
LGs (1 to 18) for the final genetic map, including 1,079 markers for progenies of the cross between GT1 and RRIM701. Black: GBS-based SNP markers; red: markers from the base map.

The SNPs identified from GBS were uniformly distributed across the genetic map (markers in black in Figure 2), resulting in improvements in group assessment and shortening of gaps compared with the base map (Figure 1). Moreover, a reduction in the number of gaps greater than 15 cM to only eight was observed, which was previously verified on 41 occasions.

### Phenotypic analysis and QTL mapping

Field-grown rubber trees were subjected to low water availability during the experimental period (Figure S1), which gave us an interesting opportunity to evaluate the performance of all genotypes under limiting conditions. In fact, stem diameter, tree height and number of whorls varied significantly throughout the experimental period and the highest rates of stem growth were found between November 2014 and June 2015, when water was less limiting (Figures S1 and S2).

BLUP values were used to perform QTL mapping, indicating the variance-covariance structures and criteria information for the selected models shown in Table 3. Table 4 shows phenotypic and genotypic values for the F1 population and each parent. Table 4 also shows the *H*^2^ for each age, and the variance components for *V_AG_* and *R_A_*.

**Table 3.** Variance-covariance structures and criteria information for the adjusted models. G_(AB,AG)_: Variance-covariance matrices for effects of replicates within block within age and genotype within age; R_(C,R, A)_: variance-covariance matrices for the column, row, and age for residual effects; df: degrees of freedom; Corh: matrix of heterogeneous variance with correlation; US: non-structured variance-covariance matrix; Ar1h: first-order autoregressive correlation matrix with heterogeneous variance; Ar1: first-order autoregressive correlation matrix without variance.

| Trait | $G_{AB}$ | $G_{AG}$ | $R_C$ | $R_R$ | $R_A$ | df | AIC | BIC | logREML |
| --- | --- | --- | --- | --- | --- | --- | --- | --- | --- |
| Tree height | US | US | Ar1 | Ar1 | US | 11 | -767.01 | -707.69 | 394.50 |
| Stem diameter | Corh | Corh | Ar1 | Ar1 | Ar1h | 26 | 22235.43 | 22407.92 | -11091.72 |
| Number of whorls | US | US | Ar1 | Ar1 | US | 11 | 1772.50 | 1831.81 | -875.25 |

**Table 4.** Summary of the predicted genotypic values and estimations of additive genetic 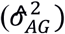 and phenotypic 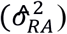 variances and heritability 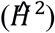 components for tree height (TH), stem diameter (SD), and number of whorls (NW) of rubber trees.

| Trait/Age | Predicted Genotypic Values | | | | | $\sigma_{AG}^2$ | $\sigma_{RA}^2$ | $\hat{H}^2$ |
| --- | --- | --- | --- | --- | --- | --- | --- | --- |
|  | F1<br>(Minimum) | F1<br>(Maximum) | F1<br>(Mean) | GT1 | RRIM701 |  |  |  |
| TH/17 months | 1.52 | 2.63 | 2.19 | 2.28 | 3.33 | 0.07 | 0.23 | 0.54 |
| TH/22 months | 2.72 | 3.85 | 3.33 | 2.06 | 2.98 | 0.11 | 0.41 | 0.51 |
| SD/12 months | 14.91 | 18.17 | 16.87 | 16.05 | 15.64 | 0.64 | 11.35 | 0.55 |
| SD/17 months | 18.72 | 23.71 | 21.76 | 20.41 | 19.84 | 1.51 | 18.14 | 0.55 |
| SD/22 months | 24.95 | 32.81 | 29.64 | 27.74 | 26.69 | 3.63 | 25.23 | 0.57 |
| SD/28 months | 31.31 | 38.66 | 36.02 | 33.96 | 32.85 | 3.27 | 39.00 | 0.54 |
| SD/35 months | 42.41 | 57.90 | 51.68 | 47.81 | 46.00 | 13.78 | 64.71 | 0.57 |
| SD/47 months | 60.15 | 82.03 | 73.69 | 68.42 | 64.62 | 29.62 | 123.66 | 0.56 |
| SD/59 months | 66.21 | 95.10 | 83.58 | 75.48 | 71.93 | 47.02 | 204.38 | 0.56 |
| NW/17 months | 5.53 | 7.69 | 6.80 | 7.00 | 6.59 | 0.22 | 1.23 | 0.45 |
| NW/22 months | 7.52 | 10.08 | 9.03 | 9.27 | 8.75 | 0.32 | 1.75 | 0.45 |

For QTL mapping analyses of the stem diameter, tree height, and number of whorls, several QTLs were identified by CIM for full-sibling progenies at different ages of growth. Using this method, 24 QTLs were identified for stem diameter, seven were identified for height, and seven were identified for the number of whorls (Table 5 and Figure 3). The QTLs were located in 12 of the 18 linkage groups, and most were in LG9 (5) for height and diameter and in LG4 (7) and LG15 (5) for all traits. A significant number of QTLs for diameter were also observed in LG17 (4). All QTLs were named with the prefix of the trait evaluated, the date (in months), and the linkage group mapped, for example, Diameter12-LG2.

**Table 5.** QTL mapping for tree height, number of whorls, and stem diameter for progenies of the cross GT1 x RRIM701. The phenotypes were evaluated at 12, 17, 22, 28, 35, 47, and 59 months. LG - linkage group; LOD - global LOD score; R^2^ - explained phenotypic variation; LOD^(2), (3), (4)^ - LOD values for additive and dominance effects.

| QTL | ID in Fig. 3 | LG | Position (cM) | LOD | R<sup>2</sup> | Additive effect for GT1 | LOD<sup>(2)</sup> | Additive effect for RRIM701 | LOD<sup>(3)</sup> | Dominance effect | LOD<sup>(4)</sup> | Segregation pattern |
| --- | --- | --- | --- | --- | --- | --- | --- | --- | --- | --- | --- | --- |
| Height17-LG4 | A | 4 | 43.41 | 6.72 | 14.07 | -0.049 | 3.256 | 0.054 | 3.965 | -0.018 | 0.496 | 1:2:1 |
| Height17-LG9 | B | 9 | 196.26 | 5.28 | 6.24 | -0.002 | 0.006 | 0.038 | 1.881 | -0.050 | 3.106 | 1:2:1 |
| Height17-LG15 | C | 15 | 108.00 | 4.23 | 7.44 | -0.004 | 0.020 | -0.039 | 2.067 | -0.049 | 2.789 | 1:2:1 |
| Height17-LG18 | D | 18 | 251.44 | 4.60 | 7.38 | -0.029 | 1.193 | -0.055 | 3.606 | -0.007 | 0.054 | 1:2:1 |
| Height22-LG5 | E | 5 | 15.00 | 5.70 | 7.18 | -0.027 | 3.782 | -0.019 | 1.734 | 0.013 | 0.828 | 1:2:1 |
| Height22-LG6 | F | 6 | 155.55 | 4.89 | 8.14 | -0.033 | 1.177 | -0.003 | 0.034 | 0.031 | 4.362 | 1:2:1 |
| Height22-LG11 | G | 11 | 189.00 | 4.37 | 8.40 | -0.003 | 0.053 | 0.026 | 3.215 | 0.017 | 1.245 | 1:2:1 |
| Whorls17-LG4 | A | 4 | 49.91 | 7.27 | 8.25 | -0.089 | 3.419 | 0.093 | 3.810 | 0.058 | 1.075 | 3:1 |
| Whorls17-LG14 | B | 14 | 26.00 | 4.13 | 1.08 | -0.067 | 1.755 | -0.134 | 2.805 | 0.012 | 0.065 | 1:2:1 |
| Whorls17-LG15 | C | 15 | 105.19 | 5.56 | 10.22 | -0.032 | 0.366 | -0.088 | 2.827 | -0.099 | 3.310 | 1:2:1 |
| Whorls22-LG4 | D | 4 | 6.04 | 8.59 | 7.67 | 0.011 | 0.853 | -0.009 | 0.531 | -0.029 | 7.036 | 1:1:1:1 |
| Whorls22-LG8 | E | 8 | 71.10 | 4.78 | 3.78 | 0.012 | 1.750 | -0.014 | 2.120 | 0.012 | 1.822 | 3:1 |
| Whorls22-LG12 | F | 12 | 152.15 | 6.26 | 5.31 | 0.009 | 1.078 | 0.017 | 3.470 | 0.011 | 1.518 | 3:1 |
| Whorls22-LG15 | G | 15 | 121.11 | 4.49 | 6.01 | -0.006 | 0.406 | -0.003 | 0.086 | -0.020 | 4.059 | 1:1 |
| Diameter12-LG2 | A | 2 | 157.93 | 4.22 | 3.80 | -0.077 | 0.619 | -0.044 | 0.224 | 0.195 | 4.017 | 1:1 |
| Diameter12-LG4 | B | 4 | 34.77 | 5.69 | 9.73 | -0.106 | 1.443 | 0.143 | 2.087 | 0.118 | 1.295 | 3:1 |
| Diameter12-LG6 | C | 6 | 170.00 | 4.72 | 8.75 | -0.140 | 2.258 | -0.057 | 0.376 | 0.164 | 2.802 | 1:2:1 |
| Diameter12-LG9 | D | 9 | 196.26 | 4.73 | 6.43 | -0.129 | 1.834 | -0.025 | 0.071 | -0.189 | 3.800 | 1:2:1 |
| Diameter12-LG17 | E | 17 | 11.05 | 4.71 | 5.13 | 0.274 | 2.179 | -0.056 | 0.442 | 0.140 | 2.471 | 1:2:1 |
| Diameter17-LG4 | F | 4 | 49.91 | 8.08 | 13.04 | -0.081 | 3.381 | 0.093 | 4.152 | 0.042 | 0.823 | 1:2:1 |
| Diameter17-LG6 | G | 6 | 163.63 | 5.09 | 6.87 | -0.042 | 0.899 | -0.040 | 0.867 | 0.087 | 4.057 | 3:1 |
| Diameter17-LG9 | H | 9 | 193.00 | 5.85 | 4.74 | -0.049 | 1.134 | -0.012 | 0.062 | -0.119 | 5.344 | 1:1:1:1 |
| Diameter17-LG17 | I | 17 | 35.00 | 5.99 | 6.58 | 0.028 | 0.406 | -0.019 | 0.205 | 0.110 | 5.068 | 1:1 |
| Diameter22-LG5 | J | 5 | 6.74 | 4.66 | 8.09 | -0.129 | 2.771 | -0.027 | 0.123 | 0.116 | 2.000 | 1:2:1 |
| Diameter22-LG9 | K | 9 | 196.26 | 6.75 | 6.61 | -0.121 | 2.593 | -0.040 | 0.274 | -0.189 | 5.480 | 1:2:1 |
| Diameter22-LG15 | L | 15 | 118.00 | 8.76 | 7.79 | -0.107 | 1.395 | -0.114 | 2.061 | -0.204 | 6.648 | 3:1 |
| Diameter28-LG5 | M | 5 | 8.00 | 8.21 | 11.40 | 0.072 | 7.216 | 0.030 | 1.224 | -0.016 | 0.342 | 1:1:1:1 |
| Diameter28-LG8 | N | 8 | 229.67 | 4.52 | 4.89 | -0.035 | 1.480 | 0.036 | 1.926 | -0.017 | 0.381 | 1:2:1 |
| Diameter28-LG11 | O | 11 | 91.00 | 6.24 | 8.08 | -0.087 | 3.214 | -0.005 | 0.041 | 0.051 | 3.438 | 1:2:1 |
| Diameter28-LG12 | P | 12 | 27.00 | 7.63 | 8.37 | -0.086 | 5.561 | 0.027 | 0.749 | 0.074 | 1.800 | 1:2:1 |
| Diameter35-LG4 | Q | 4 | 37.00 | 10.06 | 12.28 | -0.217 | 1.137 | 0.479 | 4.688 | 0.514 | 4.125 | 3:1 |
| Diameter35-LG12 | R | 12 | 136.00 | 5.74 | 3.99 | 0.190 | 0.826 | 0.013 | 0.004 | 0.490 | 4.580 | 1:1 |
| Diameter35-LG17 | S | 17 | 24.00 | 6.68 | 3.01 | 1.437 | 3.179 | 0.530 | 2.050 | 0.328 | 2.380 | 1:2:1 |
| Diameter47-LG4 | T | 4 | 28.00 | 6.29 | 10.10 | -0.261 | 1.584 | 0.510 | 3.062 | 0.222 | 0.673 | 1:2:1 |
| Diameter47-LG17 | U | 17 | 11.05 | 4.37 | 5.08 | 0.262 | 1.533 | -0.196 | 0.913 | 0.301 | 1.723 | 3:1 |
| Diameter59-LG9 | V | 9 | 144.00 | 5.61 | 9.59 | -0.316 | 3.071 | -0.173 | 0.921 | -0.337 | 2.978 | 3:1 |
| Diameter59-LG12 | W | 12 | 24.24 | 4.97 | 13.03 | 0.258 | 2.047 | 0.202 | 0.949 | -0.538 | 2.712 | 3:1 |
| Diameter59-LG15 | X | 15 | 224.00 | 4.15 | 4.65 | 0.262 | 2.058 | 0.284 | 1.915 | -0.134 | 0.438 | 1:2:1 |

**Figure 3.**
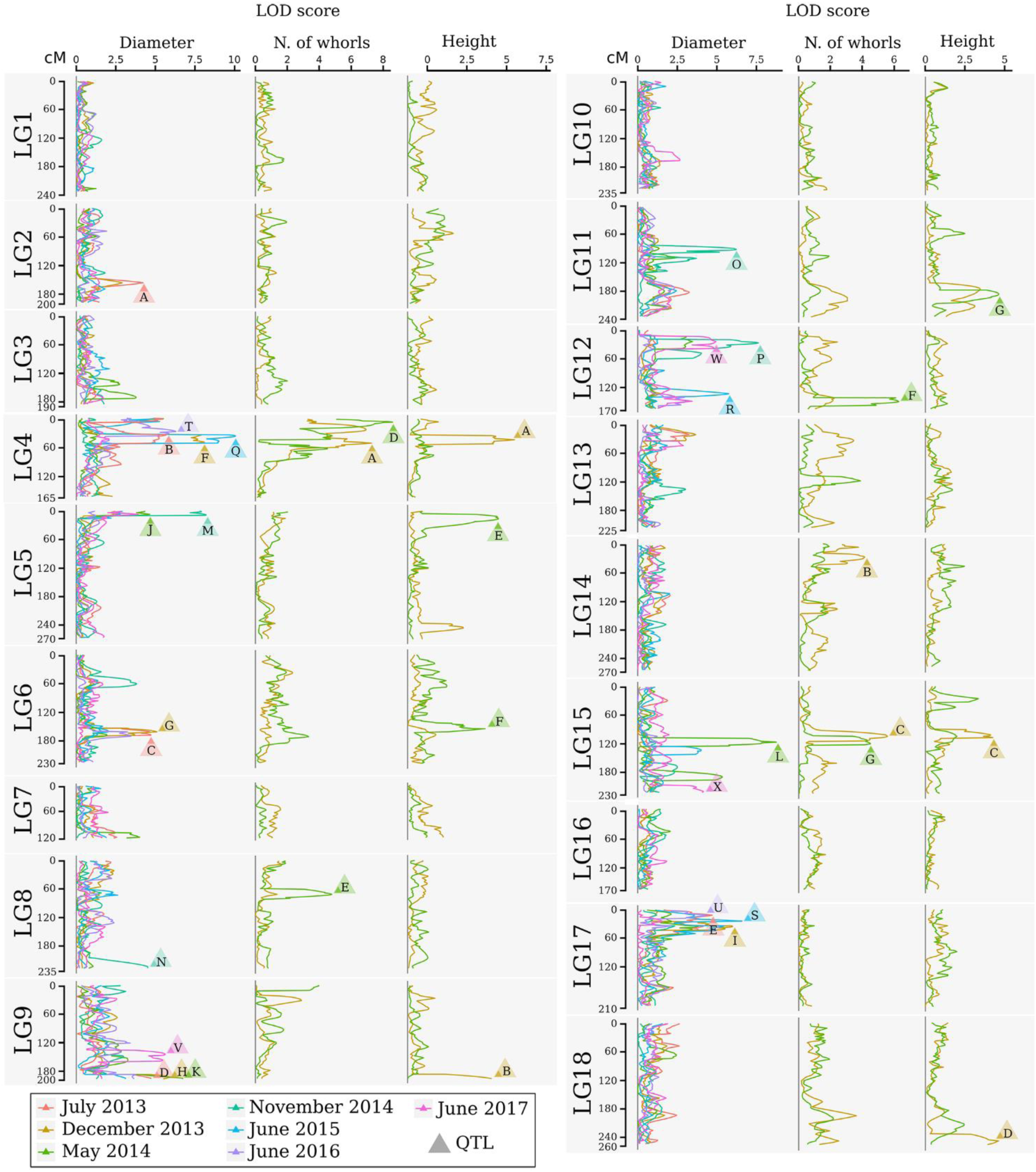
LOD profile of the QTL CIM for stem diameter, number of whorls, and tree height at multiple times for the progeny of the cross between GT1 and RRIM701. The LOD score and traits are shown on the horizontal axis. The linkage group and map position in cM are shown on the vertical axis. The colors indicate different times. Magnification allows one to visualize the genetic map and marker IDs. The markers in red also belong to the base map. LOD scores above the threshold are indicated by letters, which are described in Table 5.

From the total QTLs identified in this analysis, five were identified at 12 months, 11 at 17 months (four for height, three for the number of whorls, and four for diameter), 10 at 22 months (three for height, four for the number of whorls, and three for diameter), four at 28 months, three at 35 months, two at 47 months, and three at 59 months. The percentages of phenotypic variation (R2) explained by the QTLs ranged from 1.08% (Whorls17-LG14) to 14.07% (Height17-LG4), and the QTLs were segregated according to the ratios 1:1:1:1, 1:2:1, 3:1, and 1:1. The obtained threshold values for LOD scores using 1,000 permutations ranged from 3.94 (Diameter at 59 months) to 4.25 (Diameter at 22 months) and were used to declare the presence of QTLs.

QTLs associated with height occurred in seven LGs. QTL Height17-LG4 had the highest LOD score (6.72), whereas QTL Height17-LG15 had the lowest LOD score (4.23). Additionally, QTL Height17-LG4 had the highest proportion of phenotypic variation for this trait (14.07%), and QTL Height17-LG9 had the lowest (6.24%). For QTL Height17-LG9, Height17-LG15, and Height22-LG11, dominance and additive effects were observed for RRIM701, whereas dominance and additive effects for GT1 were observed only for QTL Height22-LG6.

For the number of whorls, QTLs were detected in five LGs. The LOD scores for these QTL ranged from 4.13 (Whorls17-LG14) to 8.59 (Whorls22-LG4), whereas the highest proportion of phenotypic variation was observed for QTL Whorls17-LG15 (10.22%). Based on the signals of the additive and dominance estimate effects, different segregation patterns for each of the QTLs were inferred. Up to three main effects (one additive effect for each parent and one dominance effect) were observed for QTL Whorls17-LG4, Whorls22-LG8, and Whorls22-LG12.

Although 24 different QTLs were mapped for the diameter trait over seven developmental stages, the most representative linkage groups mapped were LG4 (12, 17, 35 and 47 months), LG9 (12, 17, 22, and 59 months), and LG17 (12, 17, 35, and 47 months). The Diameter QTLs accounted for 3.01% (Diameter35-LG17) to 13.03% (Diameter17-LG4) of the total phenotypic variation, whereas the LOD scores ranged from 4.15 (Diameter59-LG15) to 10.06 (Diameter35-LG4). The most representative segregation patterns observed for the QTLs were of the 1:2:1 type (12 QTLs), but all types were inferred at least once. Among the 53 effects estimated for diameter, only a few were observed for RRIM701 (21 additive effects for GT1, 13 additive effects for RRIM701, and 19 dominance effects). All of the QTLs identified at 12, 22, and 35 months showed significant dominance effects.

## Discussion

The mapping population was based on a cross between two important commercial clones, GT1 and RRIM701. Figure S2 shows that the chosen parents are contrasting for the analyzed traits; their mean are outside the interquartile range. GT1 performed better in our evaluations, this can be a reflex of the environmental conditions of our trial (suboptimal region with extent drought period). GT1 is known by its climate resistance response (Cheng et al., 2015; Moreno et al., 2005; Priyadarshan et al., 2009) and suitable to be used in a study which aimed QTL mapping for drought tolerance. The construction of the mapping population was possible because GT1 is male sterile (Shearman *et al*., 2014), and there was a large experimental area with both parental clones. Using open-pollinated progenies from a mother tree of interest facilitated the development of progeny and accelerated linkage map construction and QTL detection, as in sweet cherries (Guajardo *et al*., 2015). The limited number of progenies obtained reflects the nature of the low fruit-set success of the rubber tree, varying from no success at all to a maximum of 5–10% in controlled pollination assays (Priyadarshan and Clément-Demange, 2004).

Based on these findings, the approach employed an expressive number of marker loci, with the ultimate goal of having a well-saturated map. During genetic map construction, the utilization of reliable and informative markers, such as microsatellites, might circumvent the need for phase estimation. As many markers have already been mapped in the rubber tree (Souza *et al*., 2013), use of the same markers from existing datasets is preferred to avoid confounding factors, as suggested by Hodel *et al*. (2016).

Although we observed that the proportion of polymorphic SSR markers between parents of the mapping population (48%) was consistent with that previously reported for *Hevea brasiliensis* (Lespinasse *et al*., 2000; Souza *et al*., 2013), we observed a higher proportion of EST-SSR polymorphisms than Nirapathpongporn *et al*. (2016). The number of observed polymorphic SSR markers allowed the preferable selection of highly informative markers to construct the base linkage map, e.g., the 1:1:1:1 fashion (Table 1).

Using highly informative markers together with numerous markers provided by NGS technology is advantageous. The strategy utilized herein consisted of first building a reliable base map with informative markers and then increasing its resolution with biallelic markers. To increase the reliability of markers obtained from NGS technologies, numerous GBS data filtering efforts have recently been described in several species (Peng *et al*., 2013; Ward *et al*., 2013; Guajardo *et al*., 2015; Covarrubias-Pazaran *et al*., 2016), with the objective of avoiding false polymorphism identification because of a low read depth. This objective is particularly important for large genomes containing many short repeat sequences (Fierst, 2015). To improve the genomic SNP discovery approach, a solid reference genome sequence is imperative. Thus far, a series of research studies focusing on *Hevea brasiliensis* reference genomes has been published (Rahman *et al*., 2013; Lau *et al*., 2016; Tang *et al*., 2016; Pootakham *et al*., 2017), which can be beneficial in linkage mapping studies.

Compared with other species from Malpighiales, *Hevea* species contain the largest genomes but might also have the largest repeat content (Tang *et al*., 2016). Seventy percent of the *H. brasiliensis* genome is composed of repeat elements, which, together with the lack of information at the chromosomal level, leads to difficulties with assembly (Rahman *et al*., 2013) and challenges in the identification and development of SNP assays (Souza *et al*., 2016). Moreover, differences in the estimated rubber tree genome size have been observed by microdensitometry, flow cytometry (Bennett and Smith, 1997), and 17-mer sequence estimation (Tang *et al*., 2016) approaches.

We verified the effects of inserting non-Mendelian markers in the map before including them in the base map. Segregation distortion is a common phenomenon in genetic analysis, although it might affect the estimated recombination fractions between markers (Xu, 2008). Among the GBS-based SNPs, 5,546 exhibited segregation distortion and were excluded due to the increased complexity of such a large number of markers. The analysis of such markers with segregation distortion could help identify whether one or both parents carry lethal or sublethal alleles and could aid in understanding the genetic architecture of floral development and fertility genes (Covarrubias-Pazaran *et al*., 2016).

Previous genetic map studies of *H. brasiliensis* have revealed the influences of reference genome sequences. Shortly after the first draft sequence became available for the species (Rahman *et al*., 2013), Shearman *et al*. (2015) and Pootakham *et al*. (2015) performed SNP genotyping with high-throughput platforms. Pootakham *et al*. (2015) used a read depth ≥ 6 and fewer than 50% missing data to obtain 7,345 and 6,678 SNPs in two different populations. Additionally, after using more restrictive parameters (read depth ≥ 30 and 10% missing data) and filtering uninformative markers, 2,995 and 3,124 SNPs were identified. These quantities are similar to those obtained in the present work. It is important to mention that the map construction relies on several genetic features (e.g., Mendelian segregation) present on segregating populations. Therefore, after filtering, markers were tested for these features, avoiding the presence of non-reliable markers.

The identification of SNPs in species that underwent polyploidization events, as demonstrated in *Hevea brasiliensis* (Pootakham *et al*., 2017), leads to the utilization of appropriate strategies for the selection of useful markers for linkage mapping (Clevenger *et al*., 2015). A reduction of the final number of obtained SNPs (2,998 SNPs) reflects the restrictive parameters used for marker selection. The genotyping errors and large amounts of missing data observed in NGS data (Nielsen *et al*., 2011), particularly in GBS, could have effects on the ordering of loci and the estimation of distances between two loci (Shields *et al*., 1991; Hackett and Broadfoot, 2003). Thus, these factors can critically affect the localization of regions that control quantitative traits (Doerge *et al*., 1997).

The 2,998 SNPs obtained after filtering enabled higher marker densities than those of microsatellite-based maps (i.e., Lespinasse *et al*., 2000; Le Guen *et al*., 2008; Souza *et al*., 2013). GBS methodology provides 1:2:1 and 1:1 markers (biallelic), with 1:1 as the most frequent (Table 1). The later markers are less informative because they carry information for only one parent meiosis event (Wu *et al*., 2002b). In our study, we have the advantages of both marker types, a larger number of SNPs, and greater information assessment of SSR.

Linkage group organization of the final map was based on previous studies of *H. brasiliensis* (Lespinasse *et al*., 2000; Souza *et al*., 2013) and allowed the order of the markers to be compared with those in common with other mapping populations. With 228.7 cM and 20 markers mapped, linkage group 10 was the largest linkage group obtained by Souza *et al*. (2013), and this group shared a total of 10 markers with those in our work, with the minor positioning difference between markers HB135 and HB174. A total of nine markers were positioned in the same order by Souza *et al*. (2013) in linkage group 14 at a greater density. In addition, previous genetic maps have demonstrated 18 LGs (Lespinasse *et al*., 2000; Le Guen *et al*., 2008; Pootakham *et al*., 2015; Shearman *et al*., 2015; Nirapathpongporn *et al*., 2016), corresponding to the haploid number for the species (2n = 36). The LGs obtained with 225 SSRs and 186 SNPs support the observations by Souza *et al*. (2013), demonstrating the necessity of mapping using different types of markers to account for additional LGs from incomplete coverage of the *H. brasiliensis* genome.

Although our base genetic map presented a higher mean marker density than that presented by Triwitayakorn *et al*. (2011) and Souza *et al*. (2013) when mapping 97 and 284 SSRs, respectively, marker intervals greater than 15 cM were observed on 41 occasions. As noted by Souza *et al*. (2013), such results can be explained by regions with low recombination frequency, with higher degrees of homozygosity, and with a low number of polymorphic markers, reinforcing the indispensable usage of other methodologies that provide a greater number of markers to increase resolution.

The GBS technique has promoted the discovery of thousands of polymorphic markers that are useful for the construction of high-density linkage maps. Because increasing marker densities in low-resolution regions can be beneficial for traditional mapping approaches and QTL mapping, GBS has become a valid and interesting alternative. Although GBS technology has been available for *H. brasiliensis* for several years (Pootakham *et al*., 2015; Shearman *et al*., 2015), no studies on the beneficial effects of SSR markers, such as those performed on *Brassica* species (Yang *et al*., 2016), have been performed for *Hevea*.

Our base genetic map including 225 SSRs markers was essential for avoiding problems during linkage group formation, ordering, and marker distance estimation. Inserting numerous GBS-based SNP markers reduced the average marker interval in all LGs, with a mean of 3.5 cM (Table 2). Regions with a low marker density in the base map (i.e., LG13 and LG15), with gaps up to 24.2 cM and 32.5 cM, were saturated. As in other linkage mapping studies using large-scale genotyping methodologies for *H. brasiliensis* (Pootakham *et al*., 2015; Shearman *et al*., 2015), substantial intervals (≥ 10 cM) were observed, supporting the results obtained in different plant species (Guajardo *et al*., 2015; Marubodee *et al*., 2015; Boutet *et al*., 2016; McCallum *et al*., 2016). The remaining gaps are common and even expected using GBS, mainly due to centromeric regions, which are not reached using this approach.

Our final map has a total of 1,079 markers and spans 3,779.7 cM, which reflects the molecular source and chosen mapping methodology. Comparatively, Souza et al., (2013) and Lespinasse et al., (2000) presented smaller maps, with 2688.8 cM and 2144 cM of total size, respectively. This size difference can be explained by several reasons: i) the smaller number of markers (284 and 717, respectively), cover a smaller genome region; ii) those studies used only non-NGS markers, which are likely to have fewer genotyping errors; and iii) as well as in Pootakham et al., (2015), the map from Lespinasse et al., (2000) uses strategies to reduce map size, removing improbable genotypes, such as those originating from double recombinations (Hackett and Broadfoot, 2003). However, the later strategy can also remove real recombinant events (false positives), particularly if the marker order is inaccurate (Wu et al., 2008). In our case, the consequence is to not account for those recombinant events in the further QTL mapping studies.

The constructed linkage map allowed the anchoring of 601 scaffolds (Table S3), accounting for approximately 8.1% of the current number of scaffolds available in the reference genome spanning 653.3 Mb (47.5% of the genome sequence). Therefore, although SNPs were not mapped in most parts of the scaffolds, these data constitute a representative part of the genome. Sequence contiguity of the rubber tree genome can be achieved as scaffold reviews are developed. Thus, the linkage map presented herein and the anchored scaffolds represent a foundation for future efforts.

The identification of genes underlying growth-related traits is critical for perennial crops grown in marginal areas. Tree growth is determined by different factors as cell division and expansion in apical and cambial meristems, developmental and seasonal transitions, photosynthesis, uptake and transport of nutrients and water, and responses to biotic and abiotic stresses (Grattapaglia *et al*., 2009). As consequence, tree growth is a complex phenomenon and possibly governed by many QTLs, which makes the usage of molecular markers for molecular breeding also a complex task. Our phenotypic evaluation showed values of heritability between 0.45 and 0.57 (Table 4), values higher than observed in Souza, et al., (2013). Additionally, our phenotypic model has a variance-covariance structure for the time (Ar1 or US, Table 3). The Ar1 structure allows higher correlations between close periods and, as the periods became distant, this correlation decreases. Since there were just two evaluations for the traits with the US structure, it has similar to the Ar1 pattern. These values show that a considerable part of the trait variance is related to genetic factors and provide the foundation to our QTL studies. The many growth-related QTLs identified in several LGs under suboptimal temperature and humidity conditions herein and in the study conducted by Souza *et al*. (2013) confirmed the numerous biological processes involved in *H. brasiliensis* tree growth (Figure 3). Elucidation of all the involved factors is imperative considering that rubber breeding has stagnated (Tang *et al*., 2016) and has migrated to marginal areas (Priyadarshan, 2017). The results obtained herein provide new information regarding QTL control of growth-related traits during almost five consecutive years in different seasons. Most importantly, characterizing QTLs in dry seasons can be advantageous because this strategy appears to be the most important factor for the identification of escape areas (Jaimes *et al*., 2016) and limits growth and latex production.

The climatological water balance revealed that sampling was performed in consecutive water deficit periods, although exceptions included very brief intervals with very high precipitation levels in March 2014 and January 2016 (Figure S1). In a suboptimal climate area with a lengthy dry season, Chanroj *et al*. (2017) found a single SNP associated with girth accounting for 14% of the phenotypic variance in *H. brasiliensis*. Although this was not observed herein, the QTLs identified in our work can also be used as an initial source for marker-assisted selection because seven QTLs showed major effects (phenotypic variance greater than 10%) (Table 5). These QTLs are supported by previous growth-related QTLs observed for the species under seasonal water stress (Souza *et al*., 2013). Because loss of water vapor through leaf transpiration occurs while capturing atmospheric CO2 for photosynthesis, the QTLs identified in our study provide powerful information regarding the mechanisms of water use efficiency in *H. brasiliensis*. Indeed, the search for favorable alleles for growth must continue as in the study conducted by Coupel-Ledru *et al*. (2016), which identified four QTLs related to reduced nighttime transpiration in *Vitis vinifera*.

Several periods of low water availability were noticed during the experimental period, from July 2012 to June 2017 (Figure S1). Such water deficit clearly affected stem growth, with the maximum growth rates being found when water was less limiting, i.e. between November 2014 and June 2015 (Figure S2). In fact, the trees restored their stem growth rate between the 28th and 35th months of experiment through a complex process involving the rearrangement of many metabolic pathways in a process defined as drought recovery (Luo, 2010). Such recovery capacity may play a more important role in drought resistance (Chen et al., 2016) and screening of genotypes with fast or efficient recovery of growth would be an interesting strategy from the breeding point of view.

For the stem diameter trait, we highlight the QTL Diameter35-LG17. Its genomic region is related with cadmium-induced protein AS8-like (Table S4). Because of the water deficiency, growth of roots is favored over that of leaves (Hsiao and Xu, 2000). On the other hand, root system can absorb more cadmium resulting in oxidative stress (Loix *et al*., 2017). Interesting QTLs were also observed in linkage group 5 in May (22 months) and November 2014 (28 months). Water deficit intensity in 2014 was highest and such QTLs were consistent with those previously mapped by Souza *et al*. (2013). Flanking QTL Height22-LG5, it is possible to observe a new marker developed (SNP Hb_seq_234) that has been annotated in the hydroxymethylglutaryl-CoA reductase (NADPH) pathway and mapped on scaffold0419.

Because all of the QTLs identified for stem diameter at 12, 22, and 35 months showed significant dominance effects (Table 5), their contribution appeared to play a role in phenotypic variation. Different effects, positions, and segregation patterns of QTLs detected over time must be considered with caution, particularly for *Hevea brasiliensis*, which has a long breeding cycle. Variations could be possible because different genes are involved in genotype-environment interactions or QTL regulation according to the developmental stage (Conner *et al*., 1998). In linkage group 4, QTLs with large effects were very important because the same genomic region appeared to control tree height, the number of whorls, and stem diameter.

Additionally, important QTLs in linkage group 6 have already been mapped by Souza *et al*. (2013) for both girth and tree height in a different mapping population cultivated under suboptimal growth conditions. Annotation of the sequences (Table S4) showed similarity with the *peptidyl-prolyl cis-trans isomerase FKBP15-1-like* gene, another good candidate gene worthy of further investigation. The activity of this isomerase has been shown to accelerate the process of protein folding both *in vitro* and *in vivo* (Marivet *et al*., 1994). Regulated by light in chloroplasts (Luan *et al*., 1994), it may also be regulated by stress and play an important role throughout the plant life cycle (Marivet *et al*., 1994). Therefore, further research of genes associated with abiotic stress and latex production in the rubber tree is required, as evidenced by the large number of plantations in areas with suboptimal climates.

The recent opportunity to associate high-density genetic maps with reference genomes in rubber tree is a key challenge to increase rubber production in marginal areas. The genetic map constructed herein, which included several approaches to handle thousands of markers, can be useful to improve rubber tree genome assembly, which still is fragmented into many scaffolds. The map allowed the identification of genomic regions related to growth under water stress revealing its genetic architecture. Such information can be further used as targets for genetic engineering. On the other hand, our results can be useful for breeders identify each parent alleles that contribute to increase values of the phenotypic traits. Breeders can also explore these resources to apply marker-assisted selection. Additionally, rubber tree breeding programs can benefit from the resources generated by adding these QTL effects as covariates in genomic selection models.

## Author contributions

Conceived and designed the experiments: AG, AS, PG, VG. Conducted the experiments and collected data: AC, CS, CM, IA, LHS, LMS. Analyzed the data: AC, AG, CT, JR, RA, RR. Collected phenotypic data: EJ. Wrote the manuscript: AC, AS, CT, RA. All of the authors read and approved the manuscript.

## Acknowledgements

We acknowledge Aline da Costa Lima Moraes for support with the experiments and Kaio Olímpio das Graças Dias and Jhonathan Pedroso Rigal dos Santos for helping with the phenotypic analyses.

## Supplemental figure legends

**Figure S1.** Temperature, precipitation and climatological water balance between July 2012 and August 2017 in Votuporanga, SP, Brazil. (A) Average maximum temperature, average temperature, average minimum temperature and monthly precipitation. (B) Evaluation of water excess and deficit periods. Black arrows indicate phenotypic measurements.

**Figure S2.** Boxplot of the genotypic predicted values for diameter (mm), height (m), and the number of whorls of the F1 population along the months of growth. The genotypic predicted value of the F1 parents RRIM 701 (cross) and GT1 (triangle) are also presented. Growth rates in millimeters per month (mean and standard deviation) are shown between each boxplot.

**Table S1.** SNPs developed from full-length expressed sequence tag (EST) libraries and bark transcriptome assembled de novo.

**Table S2.** Variance-covariance structures adjusted and compared based on the Akaike information criterion (AIC) (Akaike, 1974) and the Bayesian information criterion (BIC) (Schwarz, 1978). G_(AB,AG)_: variance-covariance matrices for effects of replicates within block within age and genotype within age; R_(C,R, A)_: variance-covariance matrices for column, row, and age for the residual effects; df: degrees of freedom; ID: identity matrix (homogeneous genetic variance without correlation); Diag: diagonal matrix (heterogeneous genetic variance without correlation); FA: factor analytic; Corh: matrix of heterogeneous variance with correlation; US: non-structured variance-covariance matrix; Ar1h: first-order autoregressive correlation matrix with heterogeneous variance; Ar1: first-order autoregressive correlation matrix without variance.

**Table S3.** Consensus map anchoring the final rubber tree map with 1,079 markers for the cross GT1 x RRIM701 into the scaffolds assembled by Tang *et al*. (2016).

**Table S4.** Functional annotation for genes contained within the marker interval of each identified QTL.

